# Temperature-Resolved Crystallography Reveals Rigid-Body Dominance Over Local Flexibility in B-Factors

**DOI:** 10.1101/2025.05.05.652037

**Authors:** Fernando de Sá Ribeiro, Luís Maurício T. R. Lima

## Abstract

The crystallographic B-factor (Bf), also known as the Debye-Waller factor (DWF) or temperature factor, relates to the mean square displacement of atoms (X^2^). The X^2^ may be composed of individual contributions from lattice disorder (LT), static conformational heterogeneity (H) throughout the lattice, rigid body vibration (RB), local conformational vibration (V), and zero-point atomic fluctuation (A). The Bf has been widely employed as a surrogate measure of local protein flexibility, although such relation has not been confirmed. In addition, reproducibility of the absolute B-factor is difficult to achieve, hampering the understanding of their individual contribution. Here we report the crystallographic investigation of the enzyme-ligand complex of trypsin with benzamidine from cryo to room temperature, through a 200 K range (9-point triplicate design), by crystal stabilization with hydrophobic grease. The extent of temperature-induced conformational changes showed no connection with their respective B-factors. The B-factor variation due to temperature was constant for all atoms of the system, of about 0.005 K^-1^. The results caution against interpreting absolute, normalized or zero-point B-factors as direct proxies for protein dynamics, which is further supported by structural analysis of data from independent groups with trypsin-benzamidine complexes obtained under dissimilar experimental conditions. The similar thermal dependence of B-factor for all atoms of the system suggests a major contribution of this physical variable over uniform rigid body vibration.

## Introduction

The molecular properties of proteins has been studied for over 60 years as celebrated by the seminal work of Max Perutz (Perutz *et al.* 1960) and John Kendrew (Jc *et al.* 1960) in the elucidation of the crystal structure of hemoglobin and myoglobin respectively. Conformational transitions of proteins upon physical and chemical variables has been investigated for close to a century since the pioneer work of William Astbury and Dorothy (Crowfoot) Hodgkin (Astbury & Woods 1930; Bernal & Crowfoot 1934) and the demonstration by x-ray diffraction of protein conformational transition between α-rich and β-rich structures.

Molecular dynamics lies between molecular snapshots and conformational transitions, and are essential for deeper understanding of mechanisms (Hollingsworth & Dror 2018). Dynamics is an essential part of life at all scales, from atoms to molecules, organisms and the biome. Motion can also occur at different extensions and time scales. Atomic motion can take picoseconds within a few square Angstroms (Li *et al.* 2020), while protein motion either functionally important (FIM) or biologically unimportant (BUM) (Ansari *et al.* 1985), can take nano to microseconds over tens of nanometers.

Molecular structural biology techniques such as single crystal and serial crystallography, NMR, cryoelectron microscopy (cryoEM), atomic force microscopy, molecular dynamics simulations, along with spectroscopic technics including circular dichroism and Fourier-Transformed infrared have been used in the understanding of the dynamics and conformational transitions (Lengyel *et al.* 2014). Among these techniques, crystal (single or multi) diffraction (x-ray, electron) is yet routinely used in the elucidation of polypeptide structures, and is highly reproducible as shown by protein obtained from diverse biotechnological processes and solved using data from different instruments (home sources and synchrotron) (Fávero-Retto *et al.* 2013; Liebschner *et al.* 2013). Along with coordinates, the crystallographic structures provide also a less reproductible parameter, the B-factor, which is known as temperature factor or atomic displacement parameter (Karplus & Schulz 1985; Rupp 2009; Giacovazzo *et al.* 2011; Arnold *et al.* 2012), and reflects the mean-square atomic displacement (*X*^2^) by the Debye-Waller function (DWF) as follow:

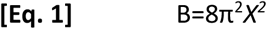

where *X*^2^ can arise from various sources, with contribution from zero-point (zero Kelvin) atomic fluctuation (*X*^2^_A_), conformational vibration (*X*^2^_v_), static conformational heterogeneity (*X*^2^_h_), rigid body vibration (*X*^2^_RB_) and crystal lattice disorders (*X*^2^_LT_) (Frauenfelder *et al.* 1979; Ringe & Petsko 1986).

The lack of correlation between B-factor and conformational changes induced by temperature challenge the popular use of B-factor as a surrogate for protein dynamics. In fact, the dissociation between B-factor and dynamics have been anticipated close to 30 years ago (Buck *et al.* 1995; Haliloglu & Bahar 1999), but is still under debate, and limited by the reproducibility of B-factor measurements.

The large variability of B-factor between measurements, crystals and setups can be empirically scaled down by its average B-factor (B_avg_) resulting in reproducible normalized B-factor (B_norm_) (Karplus & Schulz 1985; Ramos *et al.* 2022) in the form of

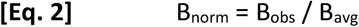

While mathematical normalization provide comparability between datasets, it is desirable to measure absolute (raw) B-factor once ensured that the crystal is stabilized, allowing the investigation of temperature as a continuous variable from cryogenic to high (above room) temperature (De Sá Ribeiro & Lima 2023) overcoming the critical temperature range of about ∼210 K which relates proteins to glasses (Ansari *et al.* 1985).

In order to obtain further insight on the linkage between B-factor and conformational transition, we used trypsin in complex with benzamidine as a case-study. Trypsin-benzamidine crystal structure has been shown to be highly reproducible at high resolution (Liebschner *et al.* 2013). In this study we investigated the temperature dependence of B-factor over temperature from 100 K to 300 K in triplicate at each temperature, using trypsin-benzamidine crystals protected with hydrocarbon grease mounted in regular nylon loops, which has the advantage of not using sophisticated systems such as capillaries and tubbing (Dunlop *et al.* 2005). We discuss our findings at light of potential correlation between observed conformational changes and B-factors.

## Material and Methods

### Material

Bovine pancreatic trypsin (Cat #SLBZ8570) and benzamidine (Cat #MKCH1700) were obtained from Merck/Sigma-Aldrich and kept at -20^o^C until use. All other reagents were of analytical grade.

## Methods

### Protein crystallography

Trypsin crystals were prepared via vapor diffusion using the sitting-drop method on Corning 3552 plates. Each drop consisted of 1.0 µL of 35 mg/mL Trypsin freshly prepared (by dissolving the protein with 5 mg/mL Benzamidine, 100 mM Hepes, and 3 mM CaCl_2_ at pH 7.0, combined with 1 µL of a precipitating agent. The drops were equilibrated against 80 µL of a reservoir solution made up of 0.2 M K_2_HPO_4_ and 20% w/v Polyethylene glycol 3,350, at a temperature of 22 ± 2 °C. Crystals suitable for diffraction emerged within 24 hours and were harvested after 48 hours. The crystals were manipulated using 20 µm nylon CryoLoops™ (Hampton Research). Each crystal was immersed in Apiezon® N hydrocarbon grease, and subsequently mounted on the goniometer in a nitrogen stream at the target temperature for data collection.

### Diffractometer setup and data acquisition

The crystals underwent X-ray diffraction and data collection using CuKα radiation with a constant exposure time. This was performed using a 30 W air-cooled µS microfocus source (Incoatec™) attached to a D8-Venture diffractometer (Bruker) at the CENABIO-UFRJ facility, operating at 50 kV and 1.1 mA. The data were captured on a Photon II detector (Bruker). Crystals were maintained under a nitrogen stream at the specified temperature with a flow rate of 1.2 L/h, regulated by a CryoStream 800 (Oxford Cryogenics). All datasets were obtained with 30-second exposures and 0.5° oscillation per image, ensuring at least 99% completeness by assuming identical Friedel pairs, and aiming for a resolution of 1.5 Å.

### Data processing and analysis

The data were collected, indexed, integrated, and scaled using Proteum3 (Bruker AXS Inc.), followed by analysis with Truncate (C.C.P.4 v7.0.071). The crystal structures were solved through molecular replacement, involving 20 cycles of rigid body search with RefMac v5.8.0238, employing PDB entry 1S0R (*Bovine Pancreatic Trypsin inhibited with Benzamidine at Atomic resolution,* at 1.02 Å). This process yielded a definitive solution for a monomer in the asymmetric unit. The initial solution underwent further refinement with 10 cycles of restrained refinement using Refmac. Real space refinement was performed by visually inspecting both the map and the model with C.O.O.T. v0.8.9.2, adjusting misplaced side chains and adding water molecules at a 1.2 σ threshold. This was followed by an additional 10 cycles of restrained refinement using Refmac. The data processing workflow was conducted using default modes in order to avoid bias. Subsequent data analysis was conducted using Superpose v1.05 from C.C.P.4. Global pairwise alignment of the trypsin-benzamidine structures was performed with ProSMART, and the analysis of crystallographic B-factor was conducted using Baverage (CCP4).

The crystallographic information on data collection and refinement statistics are informed in the **Supporting Information** (**Table S1**). Figures of crystal structures were generated using PyMOL v2.0. The atomic coordinates have been deposited in the Protein Data Bank (https://www.rcsb.org/) and the corresponding codes are provided in the **Supporting Information**. The raw images were deposited in the Integrated Resource for Reproducibility in Macromolecular Crystallography (https://www.proteindiffraction.org/).

### RCSB data analysis

A curated set of information on the orthorhombic (P2_1_2_1_2_1_) trypsin crystals with benzamidine, determined via single-crystal X-ray diffraction, was retrieved from The Research Collaboratory for Structural Bioinformatics Protein Data Bank (RCSB PDB, rcsb.org) as of May 2024 (Supporting Information). After excluding entries with incomplete data, the remaining information was plotted for the intended analysis (**Fig. S5**).

### Circular Dichroism

Thermal denaturation of Trypsin-benzamidine (5 mg/mL) was conducted in a buffer containing 0.01 mg/mL Benzamidine, 10 mM Hepes, and 300 µM CaCl_2_ at pH 7.0, utilizing a circular dichroism spectropolarimeter (JASCO® J-1000; Tokyo, Japan) equipped with a Peltier temperature control system from Jasco®. Measurements were taken using a 100 µm pathlength quartz cuvette (Uvonic Instruments, Plainview, NY). Spectra were recorded over three scans from 260 to 190 nm, with a spectral bandwidth of 0.2 nm, a scan speed of 100 nm/min, and a response time of 1 second. Corresponding experiments with blank solutions were also performed to ensure accurate background subtraction. The temperature was increased at a rate of 5 °C/min, with a 60-second equilibration period before each measurement. The resulting temperature curve was analyzed using a four-parameter sigmoidal function.

### Structural analysis from RCSB-PDB

The RCSB was searched for crystallographic structures of bovine trypsin in complex with benzamidine in space group P2_1_2_1_2_1_, resulting in 39 structures. The extracted values from these datasets included average B-factor, Wilson B-factor, Rwork, Rfree, and cell parameters. Additionally, B-factors for Cα atoms and side chains were extracted and normalized according to previous studies. Finally, using a reference structure (PDB ID 9AW0), the structures were superimposed using Superpose v1.05 from C.C.P.4 (CCP4), and the RMSD for Cα and side chains was obtained.

### Data analysis

Graphics were generated with GraphPad Prism v. 8.0.2 for Windows (GraphPad Software, San Diego, California USA, www.graphpad.com). Correlation analysis was performed using Pearson r and two-tailed distribution.

## Results

### Reproducible crystal dfWe would like to thanks the Centro Nacional ifraction at high resolution from 100 K to 300 K

The crystallographic structure of trypsin bound to benzamidine was solved at high resolution (1.5 Å) over the temperature range of 100K to 300 K in 25 K intervals, with three datasets per temperature, from a total of 27 independent crystals. The use of hydrocarbon grease for crystal protection contributed for protection against solvent loss, resulting in reproductible structures and B-factors among different crystals collected at the same temperature, and allowing collection over a wide range of temperature as a continuous variable. The crystal cell parameters showed only minor variations as a function of temperature (**Fig. S1** and **Fig. S2)**. Circular dichroism (CD) measurements of trypsin-benzamidine complex in 20 mM dibasic potassium phosphate buffer pH 7.0 confirmed the high thermal stability of the complex in solution, maintaining its structural integrity even at high temperatures (**Fig. S3**).

### Temperature-induced Conformational Changes

The alignment of trypsin structures demonstrated a high similarity between complexes at varying temperatures, as shown for backbone (**Fig. 1A**) and side chains (**Fig. 1B**), although with a progressive increase in global RMSD for both side chains (**Fig. 1C**) and Cα (**Fig. 1D**) and lack of variability in average secondary structure content (**Fig. 1E**).

**Figure 1.**
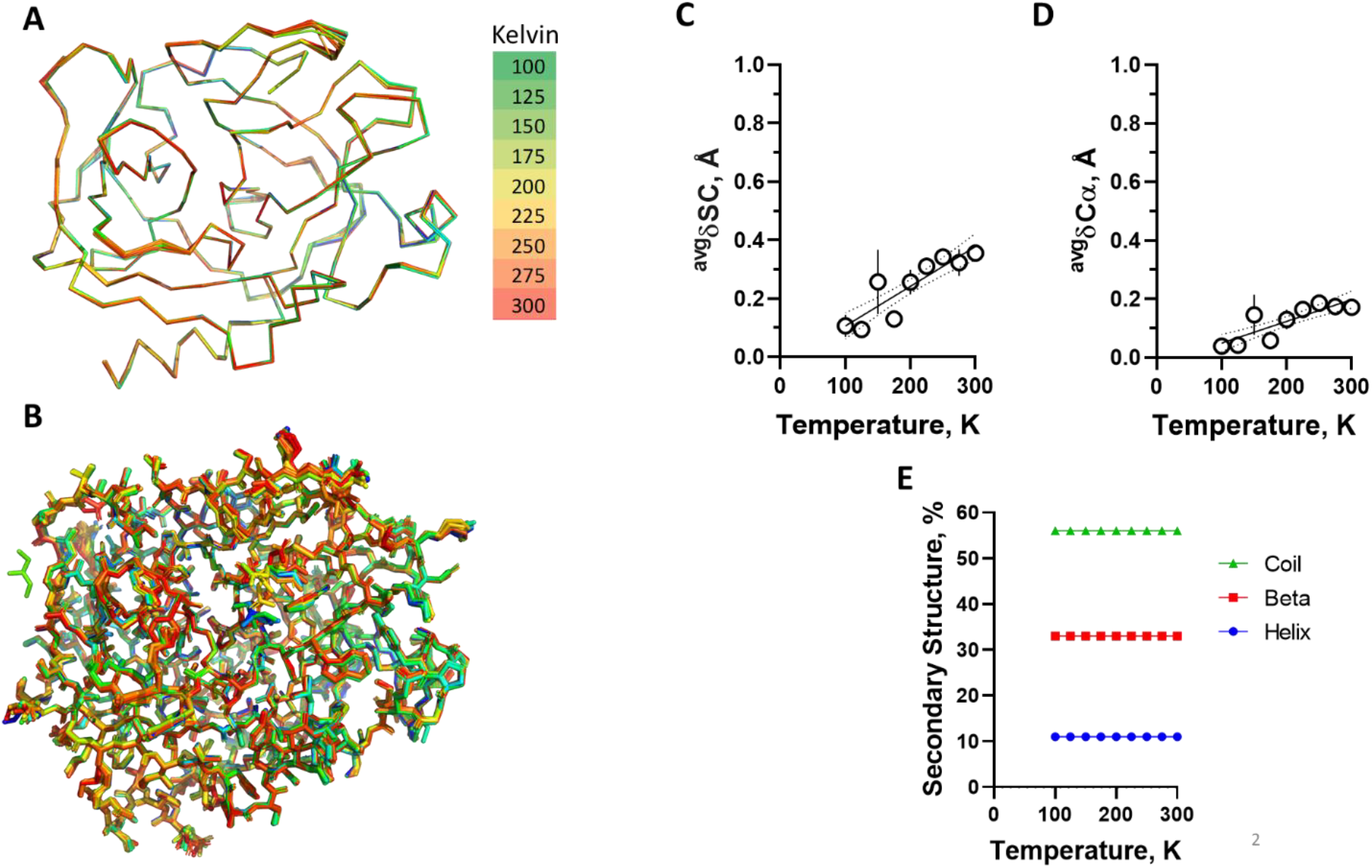
Global temperature-dependent conformational changes. Trypsin structure models solved from data collected at varying temperatures from 100 K to 300 K were aligned using the reference structure (PDB: 9AVX) and are showed for **A)** backbone and **B)** side chains. Colors represents data collection temperature as indicated. n=3 per temperature. Global (average values for residues 1-223) changes in conformation were inferred from superposition with a reference structure at 100 Kelvin. Continuous line are first order linear regression and dotted lines are 95 % confidence interval. We observed a small (< 1 Å) but progressive and significant increase in global conformational changes as inferred for distance changes in **(C)** SC (0.001358 Å.K^-1^; p≤ 0.0001) and **(D)** Cα (0.0007422 Å.K^-1^; p≤ 0.0001). The temperature changes haven’t variation in percent of secondary structure, as shown in **(E)**. Symbol is average and bar is standard deviation (n=3).

From the crystal structures, local conformational changes induced by temperature were analyzed for both side chains (**Fig. 2A**) and Cα (**Fig. 3A**), with the latter showing a smaller amplitude of displacement. However, both chains exhibited non-uniform responsiveness to temperature, suggesting that temperature influences chain displacement in a localized manner.

**Figure 2.**
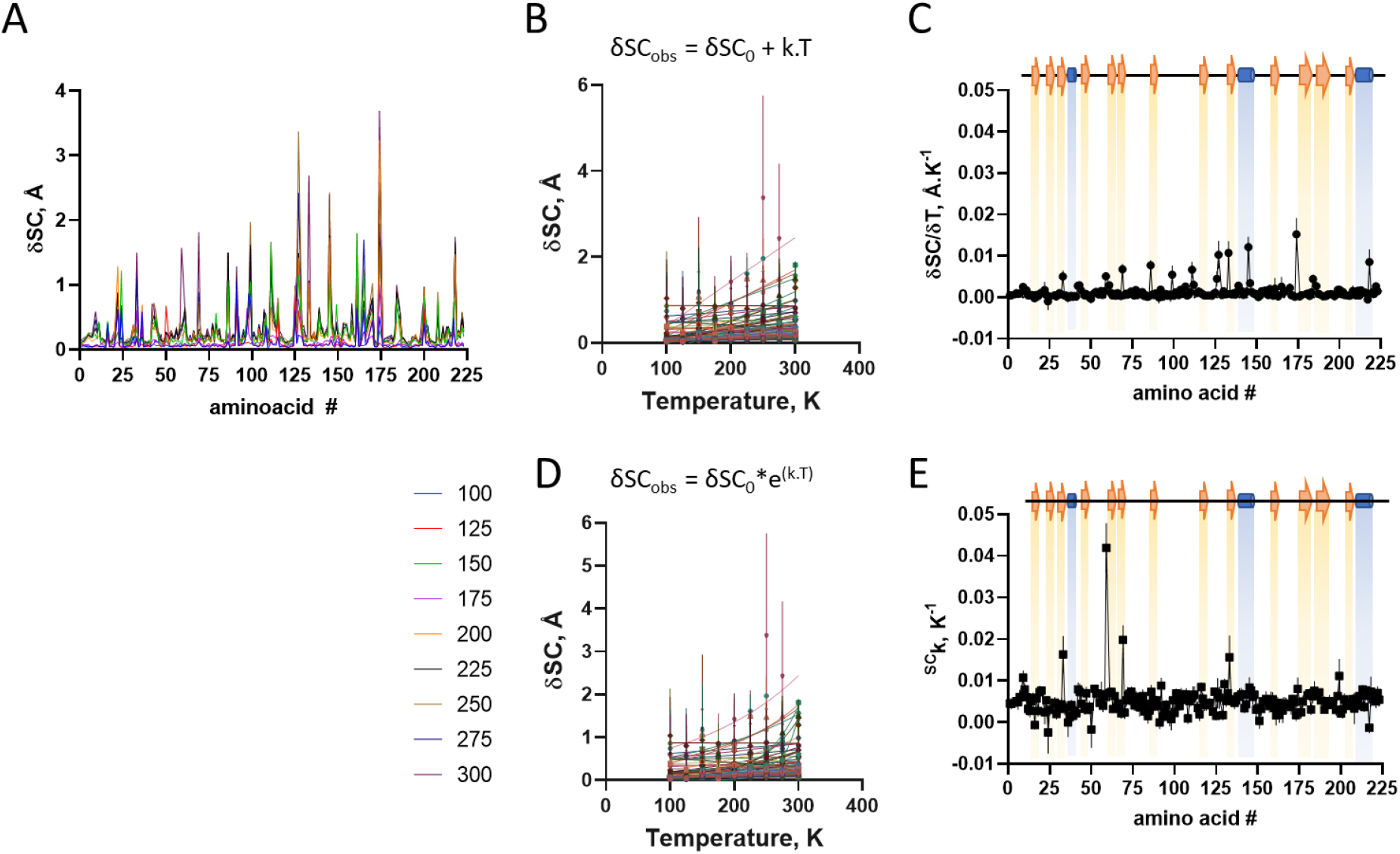
Temperature induced conformational changes in trypsin side chain. Structures were aligned with a reference structure at 100 K using *Superpose* (CCP4). **A)** Changes in SC distances along protein sequence at varying temperatures (different colors). **B)** Changes in SC distances as a function of temperature for each amino acid residue (different colors) protein sequence at varying temperatures. Lines are first order linear regression, from which were obtained their thermal-dependence of conformational change (δSC; angular coefficient of linear regression). **C)** Distribution of thermal-dependent conformational change (δSC) along protein sequence. **D)** Changes in Cα distances as a function of temperature for each amino acid residue (different colors) protein sequence at varying temperatures. Lines are exponential non-linear regression, from which were obtained their exponential thermal constant (^δSC^k). **E)** Distribution of exponential thermal constant (^δSC^k) of conformational change along protein sequence. Symbol is average and bar is standard deviation (n=3).

**Figure 3.**
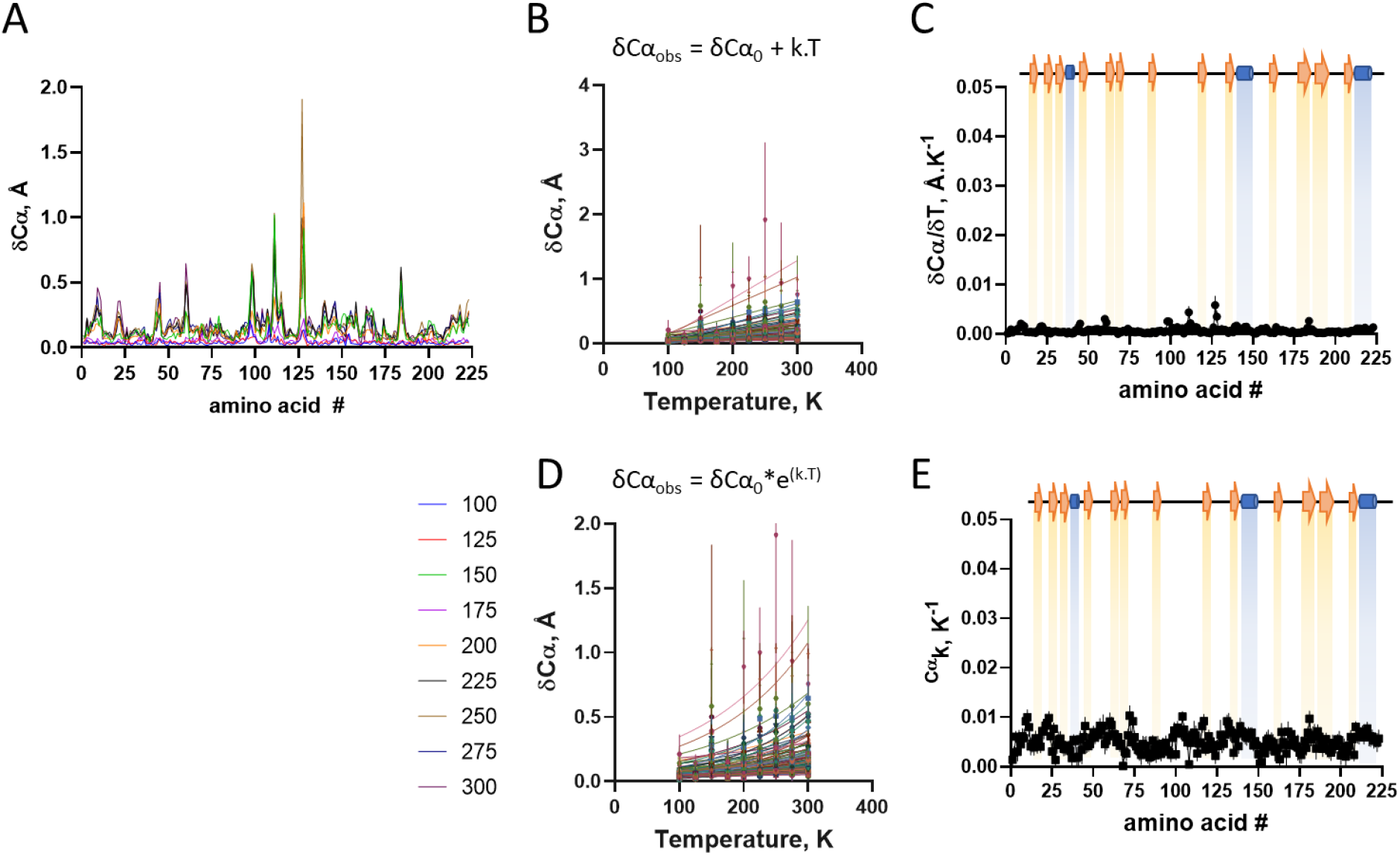
Temperature induced conformational changes in trypsin Cα. Structures were aligned with a reference structure at 100 K using Superpose (CCP4). **A)** Changes in Cα distances along protein sequence at varying temperatures (different colors). **B)** Changes in Cα distances as a function of temperature for each amino acid residue (different colors) protein sequence at varying temperatures. Lines are first order linear regression, from which were obtained their thermal-dependence of conformational change (δCα; angular coefficient of linear regression). **C)** Distribution of thermal-dependent conformational change (δCα) along protein sequence. **D)** Changes in Cα distances as a function of temperature for each amino acid residue (different colors) protein sequence at varying temperatures. Lines are exponential non-linear regression, from which were obtained their exponential thermal constant (δCαk). **E)** Distribution of exponential thermal constant (δCαk) of conformational change along protein sequence. Symbol is average and bar is standard deviation (n=3).

The temperature-induced changes were analyzed by model-free regression, using first-order linear regression **[Eq. 3]** and non-linear regression with a single exponential **[Eq. 4]** for Cα (**Fig. 2B-D**) and side chains (**Fig. 3B-D**):

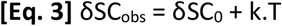

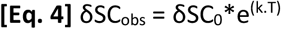

The temperature-dependence variation constants were plotted as a function of the polypeptide chain (**Fig. 2C-E** and **Fig. 3C-E**; **Fig. S4**), and are shown to vary only marginally through the polypeptide chain (**Fig. 2C** and **Fig. 3C**, respectively), suggesting the lack of major local propensity in conformational changes in response to temperature

### Global changes in B-factor by temperature

We analyzed the average changes in B-factor as a function of temperature. A positive exponential profile with temperature was observed for the B-factor derived from the Wilson plot (**Fig. 4A**), and the average B-factor from the side chains (**Fig. 4B**) and from the Cα (**Fig. 4C**). The thermal dependence of these observed average B-factors was adjusted using the single exponential function **[Eq. 5]**:

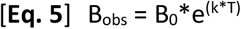

**Figure 4.**
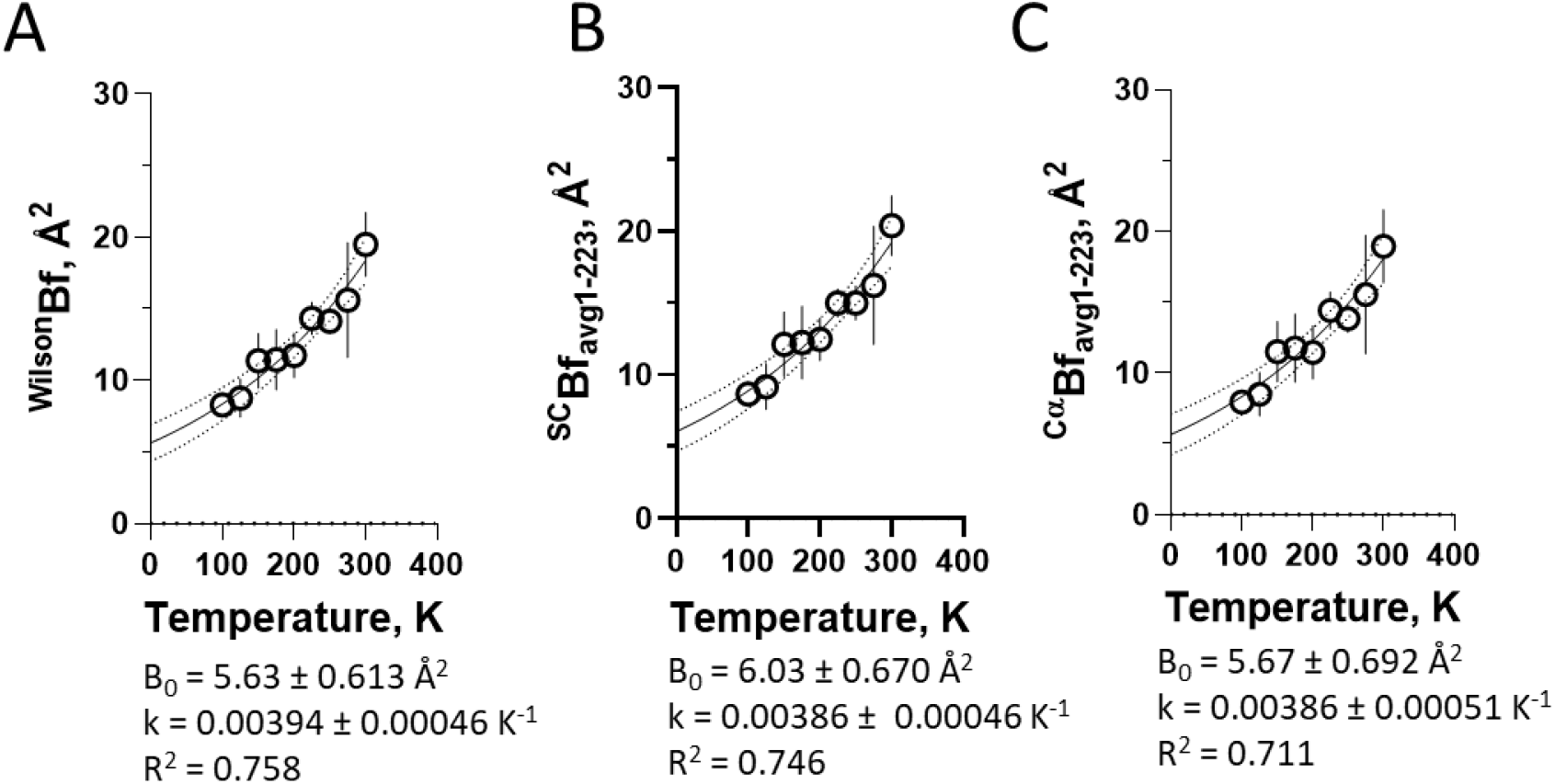
Temperature-dependent changes in B-factor (B). Overall changes in B-factor as a function of temperature for **A)** Wilson plot (Pearson r correlation coefficient = 0.9597, with 95% CI 0.8151 to 0.9917), p-value two-tailed <0.0001), **B)** Side-chains (Pearson r correlation coefficient = 0.9599, with 95% CI 0.8160 to 0.9918), p-value two-tailed <0.0001) and **C)** Cα (Pearson r correlation coefficient = 0.9577, with 95% CI 0.8065 to 0.9913), p-value two-tailed <0.0001). Values corresponds to the extrapolated B factor at zero Kelvin (B_0_) and the thermal coefficient k, as inferred from single exponential (**B_obs_ = B_0_*e^(k.T)^**) non-linear regression of their respective panels (solid lines; dotted lines are 95% confidence interval). Symbol is average and bar is standard deviation (n=3).

where **B_obs_**is the refined B-factor, **B_0_**is the B-factor extrapolated to zero Kelvin from regression, **k** is the thermal constant, and T is the data collection temperature (in Kelvin). This adjustment resulted in close average thermal factors of about ^avg^k = 0.003944 K^-1^ and in ^avg^B_0_ of about 5.63 Å^2^ for the B-factor dependence on temperature in the model-independent data from Wilson plot and from the protein models (side chains and Cα).

Atomic B-factor changes by temperature

The B-factor is variable through the atoms of the system. The B-factor varies broadly along the polypeptide sequence as seen for side chains (**Fig. 5A**) and Cα (**Fig. 6A**). The B-factor distribution pattern through the polypeptide chain is similar for all temperatures although at different levels. The variation of the B-factor for a given atom (eg, Cα, **Fig. 5B**) or group of atoms (ex, side chains, **Fig. 6B**) as a function of temperature shows an exponential rise pattern, and the exponential constant show only minor variability for Cα (**Fig. 6C**) and side chain (**Fig. 5C**) of about 0.005 K^-1^, suggesting a model-free independent behavior of changes in atomic displacement parameter as a function of temperature.

**Figure 5.**
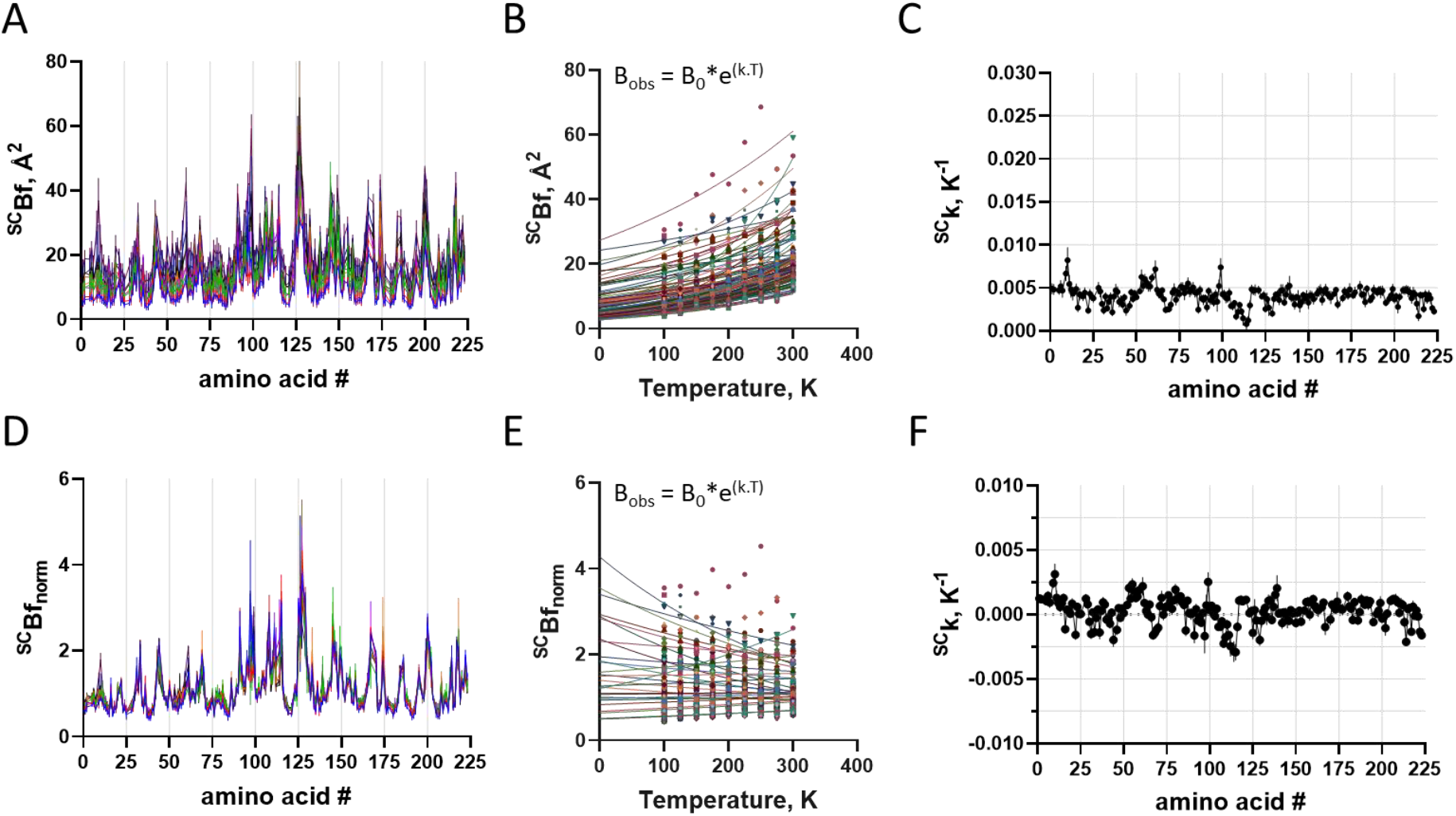
Temperature-dependent changes in side-chain B-factor. **A)** B-factor distribution for side chain at varying temperatures (different colors). **B)** Changes in raw B-factor as a function of temperature for each amino acid side chain (different colors). Lines are exponential non-linear regression, from which were obtained the raw B_0_ and the thermal constant k; **C)** Distribution of raw k along protein sequence. Pearson r correlation coefficient -0.1413 95% CI: -0.2753 to -0.001847), P (two-tailed) = 0.0471; **D)** Normalized B-factor (B_norm_) distribution for side chain at varying temperatures (different colors); **E)** Changes in B_norm_ as a function of temperature for each amino acid side chain (different colors). Lines are exponential non-linear regression, from which was obtained the thermal constant ^norm^k; **F)** Distribution of ^norm^k along protein sequence. Pearson r correlation coefficient -0.09924 (95% CI: -0.2354 to 0.04076), P (two-tailed) = 0.1642; Symbol is average and bar is standard deviation (n=3).

**Figure 6.**
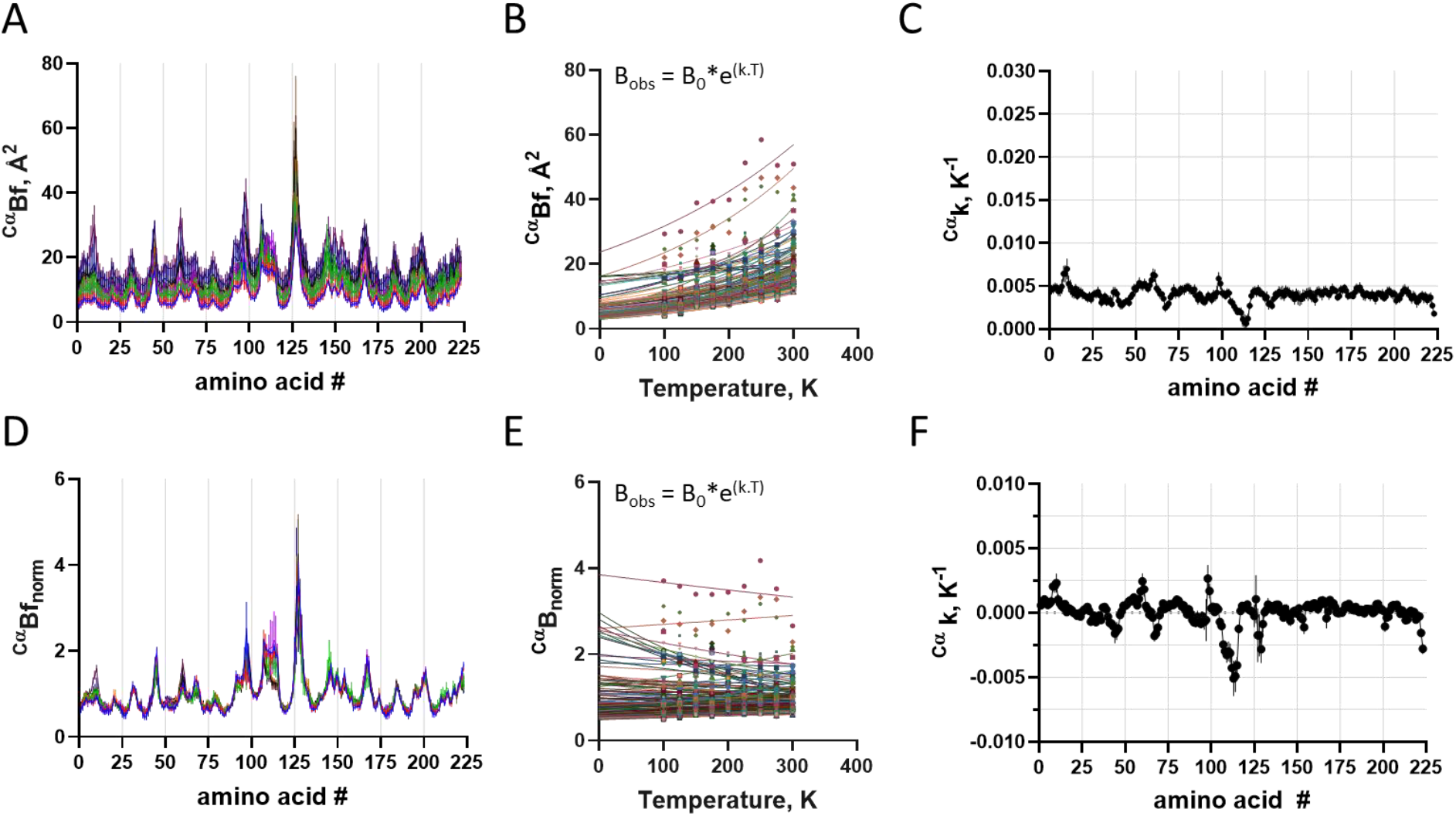
Temperature-dependent changes in Cα B-factor. **A)** B-factor distribution for Cα at varying temperatures (different colors); **B)** Changes in raw B-factor as a function of temperature for each amino acid Cα (different colors). Lines are exponential non-linear regression, from which were obtained the raw B0 and the thermal constant k; **C)** Distribution of raw k along protein sequence. Pearson r correlation coefficient -0.1819 (95% CI: - 0.3059 to -0.05173), P (two-tailed) = 0.0065; **D)** Normalized B-factor (B_norm_) distribution for Cα at varying temperatures (different colors); **E)** Changes in B_norm_ as a function of temperature for each amino acid Cα (different colors). Lines are exponential non-linear regression, from which was obtained the thermal constant ^norm^k; **F)** Distribution of ^norm^k along protein sequence. Pearson r correlation coefficient -0.09912 (95% CI: -0.2275 to 0.03268), P (two-tailed) = 0.1401. Symbol is average and bar is standard deviation (n=3).

The normalization of the polypeptide B-factor allows better understanding of local fluctuation. The normalized B-factors for Cα atoms (**Fig. 6D**) and side chains (**Fig. 5D**) resulted in similar distribution pattern at all temperatures for crystal structure elucidation between 100K to 300K. Adjusting the temperature-dependence of the normalized B-factor with a single exponential non-linear regression function according to **[Eq. 5]** for Cα (**Fig. 6E**) and side chain (**Fig. 5E**) shows only minor variation, with constant ^norm^K fluctuating around zero for both Cα (**Fig. 6F**) and side chain (**Fig. 5F**), suggesting a lack of local variation of B-factor as a protein structure feature. A minor variation may be depicted in the amino acid region 108-112, although with no statistical significance (**Fig. S5**) which may be attributed to crystallographic contact (**Fig. S6**).

### Correlation Between Conformational Change and B-Factor

A cross-correlation analysis between B-factor and conformational changes was performed. The variation of the extrapolated zero-point B-factor (zero Kelvin, B_0_) for atoms from side-chains and Cα were analyzed as a function of temperature-dependent conformational change (**Fig. 7**).

**Figure 7.**
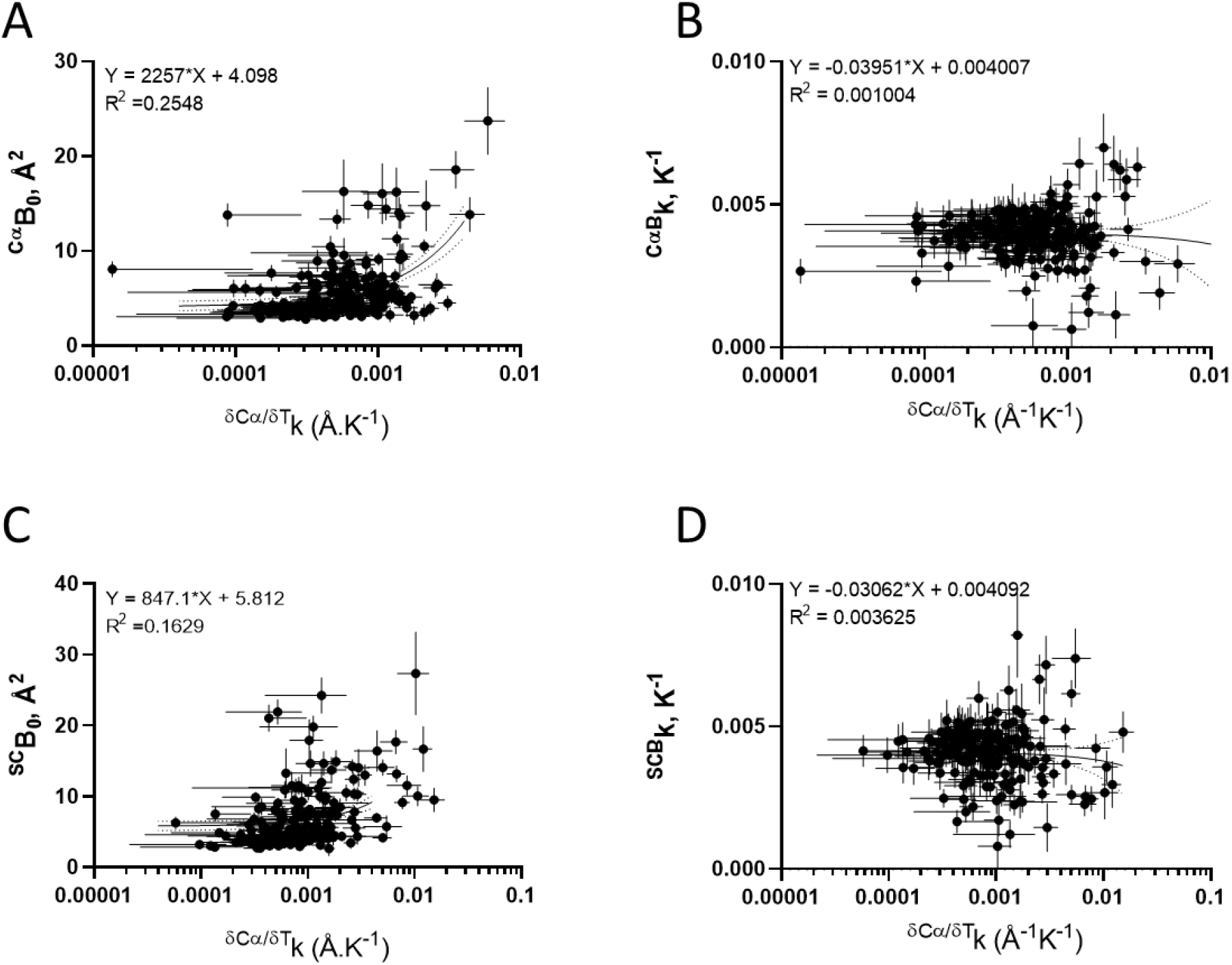
Correlation between B-factor and conformational changes. Correlation between temperature-induced conformational changes constant (δSC/δTk) and the extrapolated zero-point B-factor (B_0_, **A** and **C**) or the thermal B-factor dependence constant (Bk, **B** and **D**) for Cα (**A** and **B**) and side chains (SC; **C** and **D**). Lines correspond to first-order linear regression (continuous) and 95 % confidence interval (dotted). Pearson r correlation coefficient (log x scale): (A) r = 0.3576 (95 % CI: 0.2374 to 0.4670), P (two-tailed) <0.0001; (B) r = 0.0228 (95% CI: -0.1089 to 0.1538), P (two-tailed) = 0.7344; (C) r = 0.4553 (95% CI: 0.3354 to 0.5608), P (two-tailed) <0.0001; (D) r = -0.0761 (95% CI: -0.2154 to 0.06622), P (two-tailed) = 0.2941.

The extrapolated zero-point B-factor (B_0_, **Fig. 7A** and **Fig. 7C**) and the thermal B-factor dependence constant (^B^k, **Fig. 7B** and **Fig. 7D**) is distributed along a broad range of temperature-induced conformational changes constant (^δSC/δT^k) and lacking strong correlation between them (Pearson r <|0.5|), indicating that the minor local variation found in B-factor does not associate with the conformational changes induced by temperature.

### Influence of Temperature on the B-Factor of Non-Protein Atoms

The effects of temperature on non-protein atoms were also analyzed. The crystal structure solved at 1.5 Å allowed determining a large set of water molecules with a broad distribution of B-factors, ranging from 5 Å^2^ to 60 Å^2^ (**Fig. 8A**). The number of crystallographic water molecules decrease as a function of temperature (**Fig. 8B**), while their respective B-factor increases (**Fig. 8C**), similar to other non-protein atoms such as calcium (**Fig. 8E**) and benzamidine (**Fig. 8F**). The variation in non-protein B-factor as a function of temperature was adjusted using a single exponential non-linear regression function **[Eq. 5]**, revealing a constant fluctuating at about 0.005 K^-1^ which is in the same magnitude range for all other protein atoms from side chain and Cα. These data reveal a systematic model-free pattern in B-factor change for all atoms of the system as a function of temperature regardless of the nature of the atom.

**Figure 8.**
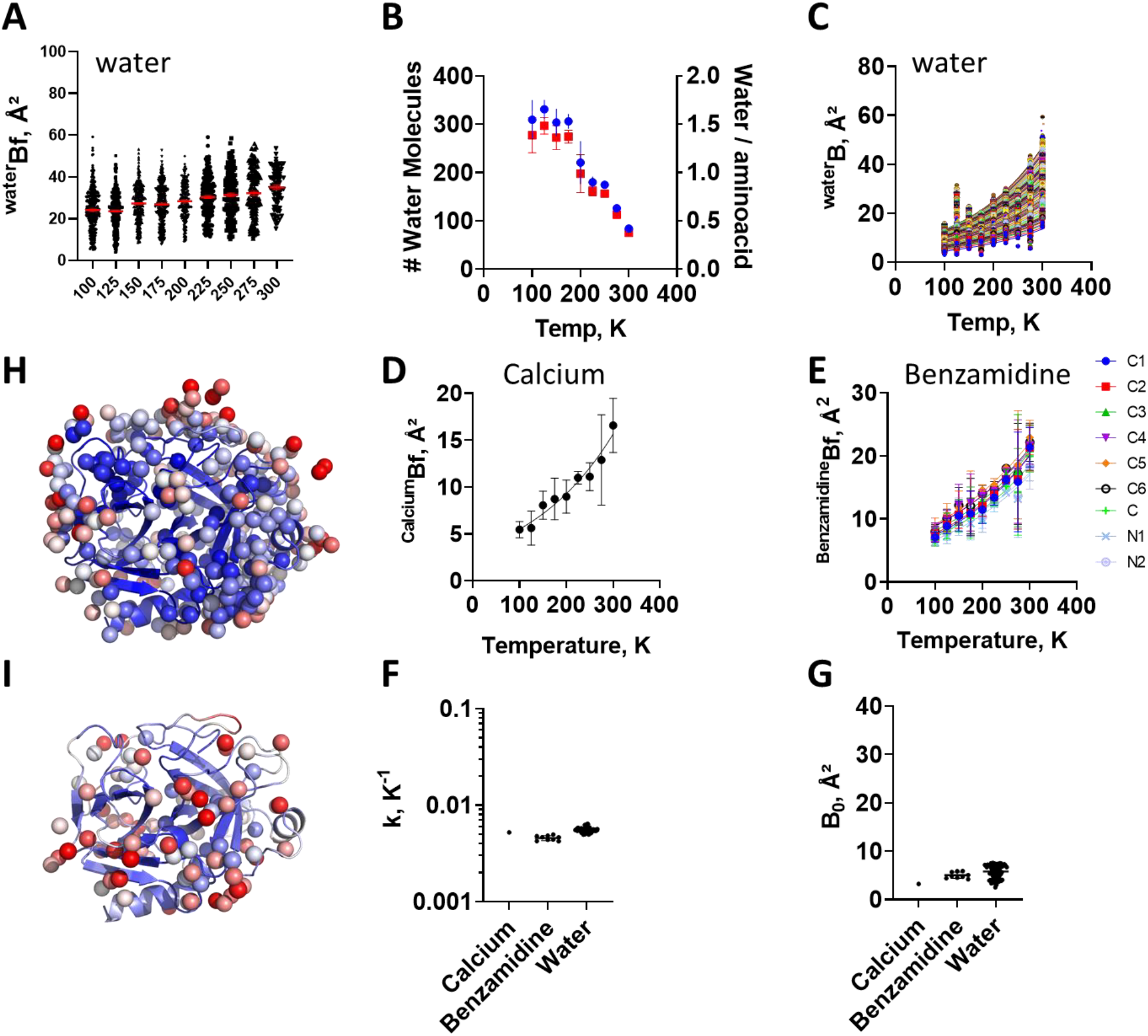
B-factor of non-protein atoms as a function of temperature. **A)** B-factor of the oxygen atoms of the water present in the structure as a function of temperature. **B)** Number of water molecules per structure and the water/amino acid quotient. **C)** B-factor of all oxygen atoms in conserved water as a function of temperature. Crystallographic structure represented by cartoon colored according to B-factor (Blue-white-red, min 5Å^2^, max 40 Å^2^), showing water molecules as spheres, Thermal dependence of B-factor for **D)** calcium ions, and **E)** Benzamidine atoms. Lines are exponential non-linear regression, from which were obtained the thermal constant k **(F)** and the raw B_0_ **(G)** (k = δB/δT). **H)** PDB:9AVX and **I)** PDB:9AWU showing the loss of molecules at 300K. Bars are average and standard error (n = 3).

### Analysis of Independent Structures from RCSB

The general effect of temperature and B-factor reported here searched from independent research groups, in other trypsin-benzamidine structures (**Fig. S7, Fig. S8**; **Supporting Material – Excel file with RCSB Custom Report**), varying in resolution (from 0.75 to 2.4 Å), pH, crystallization conditions, data acquisition and processing workflows among other differences which bring adequate high methodologic variance for this analysis.

A total of 39 independent crystal structures of bovine trypsin-benzamidine in P2_1_2_1_2_1_ were found in the RCSB. These structures were collected either at cryogenic or room temperature range, revealing a gap between these two temperature ranges (**Fig. S7A**). This gap may be due to individual interest in each of these temperature ranges, or due to difficulties to use temperature as continuous variable in this range. These crystal lattices show close cell parameters, although four of them fall outside the range (**Fig. S7B, Fig. S7C**). The changes in cell parameters showed a positive correlation between **b** vs **a** (**Fig. S7E**) and **c** vs **a** (**Fig. S7F**). These structures show close correlation between data (from Wilson plot of integrated diffraction intensities) and model (final refined structure) B-factors (**Fig. S7H**). These structures show good correlation between Rw and Rf, which minor correlation with Wilson B-factor.

The B-factor analysis of these structures reported from independent groups showed large variation in the B-factors distribution along the polypeptide chain and conformational change (**Fig. 9**), both for Cα (**Fig. 9A** and **Fig. 9C)** and side chains (**Fig. 9D** and **Fig. 9F**). However, upon B-factor normalization their distribution pattern along the polypeptide chain become in close similarity (**Fig. 9B, Fig. 9E**), indicating that the variations in conformation are not followed by changes in B-factor. Instead, a dissociation between these two parameters is found.

**Figure 9.**
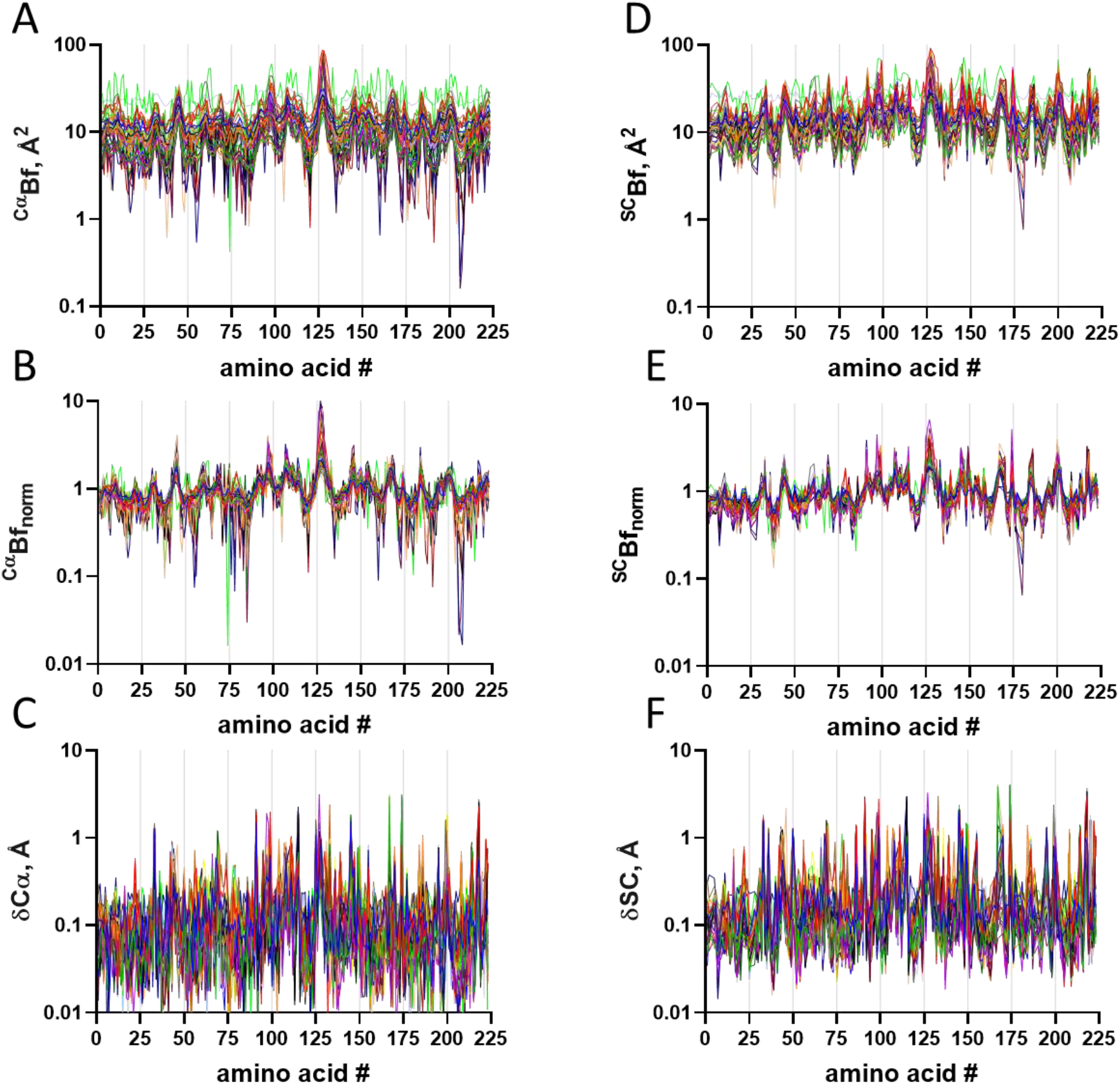
Orthorhombic trypsin with benzamidine in the RCSB. Data from the RCSB (access May, 2024) for trypsin in P2_1_2_1_2_1_. **A)** Distribution of the Cα Raw b-factor for each PDB structure. **B)** Distribution of the Cα Norm b-factor for each PDB structure. **C)** Distribution of the Cα RMSD for each PDB structure. **D)** Distribution of the Side Chain Raw b-factor for each PDB structure. **E)** Distribution of the Side Chain Norm b-factor for each PDB structure. **F)** Distribution of the Side Chain RMSD for each PDB structure.

Together, our present data provide evidences that crystal stabilization allow reproducible measurements in a broad temperature range from cryo to room temperature. Moreover, from present data and independent data from other research groups, we have found that while physical and chemical variables (including temperature, pH crystallization conditions) can result in modifications in the crystallographic B-factor, this parameter does not seem to correlate with conformational changes.

## Supporting information

Crystallographyc Information

Trypsin-Benzamidine data from RCSB

## Abbreviations

CD: circular dichroism.

## Discussion

In this study, we report a structural analysis of trypsin in complex with benzamidine solved at varying temperatures over a 200 K interval, providing insights into the protein conformational diversity and its relationship with the crystallographic B-factor. Local conformational changes induced by temperature follow a non-linear pattern, which does not correlate with their respective atomic B-factors and temperature-dependent changes. While the B-factors provide insight on the mean-square displacement of atoms, they do not directly correlate with local conformational changes. Such dissociation between conformation and B-factors is also found in other structural studies, emphasizing the complex nature of conformational flexibility and also the contributors to the average B-factor.

The three ‘Rs’ that comprise high-quality scientific research are known as rigor, reproducibility and robustness (Garraway 2017). Achieving reproducibility in B-factor measurements has proven challenging, hindering the understanding of each factor’s specific contribution. To address this issue, we first proposed a mathematical normalization (Ramos *et al.* 2022), which revealed the high reproducibility of B-factors between crystals collected at the same temperature by different instruments, analysts, and data processing workflows, and can assist ensemble refinement (Du *et al.* 2023; Beton *et al.* 2024; Leitner *et al.* 2025).

Our normalization method, already incorporated into practice (Du *et al.* 2023; Mlynek *et al.* 2024), did not solve the reproducibility issue regarding the raw B-factor and did not explain the differences between similar crystals in the same of different instruments. Recently, we demonstrated the use of hydrocarbon grease to protect lysozyme protein crystals for data collection over a broad temperature range (cryogenic to 325K) at high resolution (1.5 Å) in triplicate (De Sá Ribeiro & Lima 2023). Protective measures against dehydration for single-temperature data collection have been shown using specific devices (Dunlop *et al.* 2005) or embedding the crystals into hydrophobic chemicals, such as lipid cubic phase (LCP) (Ihara *et al.* 2020), oils (Hope 1988; Hope *et al.* 1989; Warkentin *et al.* 2012; Huang *et al.* 2024), mineral oil (Sugahara *et al.* 2015), shortening (Nam 2020a), and fat (Nam 2020b, 2022). This technique shows potential for reproducible measurements using temperature as a physical variable, which allowed us to find a lack of correlation between B-factor changes and temperature-induced conformational transitions using both normalized and raw B-factors (De Sá Ribeiro & Lima 2023).

Temperature effects on non-protein atoms were also investigated. Initially, a broad distribution of B-factors for water molecules was observed. While the number of crystallographic water molecules decreased with temperature, the B-factors of such remaining water molecules increased uniformly as a function of temperature, as also observed for protein atoms, indicating consistent behavior across different atom types. Similarly, other non-protein atoms such as calcium and the benzamidine molecule exhibited similar effects on B-factor as a function of temperature, providing evidence that the temperature-induced changes in B-factors are uniform for all atoms of the system. The thermal expansivity constants for non-protein atoms were comparable, reinforcing the homogeneous impact of temperature across various atomic components analyzed in the study.

The analysis of RCSB entries for trypsin-benzamidine crystal structures from other groups revealed heterogeneous conformations while similar in the normalized B-factors. This result has multiple interpretations. First, a demonstration of the importance of elucidating crystal structures under varying chemical and physical conditions in order to explore the conformational space that can be populated. Second, the robustness of B-factor normalization for comparative structural analysis. Finally, the lack of correlation between changes in conformation and B-factor.

In our crystallographic examination of the temperature effect on trypsin-benzamidine enzyme-ligand complex, we found no discernible correlation between temperature-induced conformational changes and B-factors. The B-factor variation due to temperature remained consistent across all atoms within the system. These findings suggest a detachment between absolute B-factor values and conformational plasticity within this particular system. The similar thermal dependence of B-factors for all atoms suggests a major contribution from uniform rigid-body vibration of the whole system rather than localized flexibility.

The pioneering work of Fraudenfelder, Petsko and Tsernoglou with crystal structures of myoglobin solved from data collected at temperatures ranging from 220 K to 300 K demonstrated the thermal dependence of B-factor on temperature, although not allowing separation between the vibrational (*X*^2^_v_) and rigid-body (*X*^2^_RB_) terms (Frauenfelder *et al.* 1979; Ringe & Petsko 1986). In our present work, we could determine that the rigid-body vibration contributes with a single exponential constant of 0.005 K^-1^ (**Eq 5**), while the zero-point B-factor (B_0_) may show contributions from conformational substates, lattice contacts and disorder, and atomic and local (conformational) vibration. In this context, the lack of correlation between conformational changes and zero-point B-factor suggests that either absolute or rescaled B-factor does not seem to be the best approach for inferring dynamics and conformational plasticity.

The interpretation of crystallographic B-factor as an indicator of local protein flexibility is tempting, and remains pervasive in structural biology literature (Sun *et al.* 2019). However, this conceptual link—between B-factor amplitude and conformational mobility — has not been supported by experimental validation. Instead, the dissociation between these two variables has originally been demonstrated by Prof. Christopher Dobson and colleagues, using lysozyme and dynamic measurements by NMR (Buck *et al.* 1995). Using primary crystallographic data of lysozyme and insulin from our group and secondary, independent data, from other groups we have additionally shown that variations in B-factor do not correlate with local conformational transitions (Ramos *et al.* 2022; De Sá Ribeiro & Lima 2023). In the present work, using trypsin-benzamidine as an independent structural model, we confirmed and extended these findings on the dissociation of B-factor and conformational plasticity, while adding evidence for the uniform, all-atoms variation in B-factor as a function of temperature, most likely due to whole-system rigid body motion. In conjunction, all these independent data from distinct proteins indicate that B-factors, either the raw values or rescaled, may not be the best surrogates for protein dynamics. The generality of these observations across structurally unrelated proteins and under distinct crystallographic conditions argues for a reevaluation of the common assumptions linking B-factors to functional flexibility. Drawing the folding landscape of proteins, their dynamics and conformational plasticity may be better accessed by techniques such as multiple conformers from serial crystallography, multiple single-crystal structures from independent datasets, use of chemical (e.g.: pH, salts) and/or physical (temperature, pressure) variables, and molecular dynamics by simulation, normal modes and/or NMR (De Magalhães *et al.* 2023; de Araujo *et al.* 2025). Solving multiple structures from independent datasets and using varying techniques may provide access to the understanding of conformational plasticity and polymorphism, and the understanding of the folding landscape of proteins.

## Conclusions

Our data establish that hydrophobic embedding stabilizes crystals for reproducible B-factor analysis across temperature ranges from cryogenic to room and higher temperatures, while revealing that B-factors predominantly reflect global vibrations rather than local dynamics. This necessitates reevaluating their use as flexibility proxies. We advocate for integrative approaches combining ensemble crystallography, MD simulations, and thermodynamic analyses to decipher conformational landscapes.

## Acknowledgments

We would like to thanks the Centro Nacional de Biologia Estrutural e Bioimagem (CENABIO, UFRJ) for the access to the microscopy facility. We would like to acknowledge the Ministério da Ciência, Tecnologia e Inovação (MCTI) and the Centro Nacional de Biologia Estrutural e Bioimagem (CENABIO) for the support with the X-ray diffraction facility (D8-Venture), and the National Institute of Science and Technology (INCT) for Structural Biology and Bioimaging / INCT-CNPq Program).

## Data availability

The experimental information and data supporting the findings of this study are available within the paper and the indicated data repository, under PDB ID listed in Table S1 (9AVX, 9AVY, 9AVZ, 9AW0, 9AW1, 9AW2, 9AW4, 9AW8, 9AW9, 9AWA, 9AWB, 9AWC, 9AWD, 9AWF, 9AWG, 9AWH, 9AWI, 9AWL, 9AWM, 9AWN, 9AWO, 9AWP, 9AWQ, 9AWR, 9AWS, 9AWU, 9AWV, and 9AWZ). Further information is available from the corresponding authors upon reasonable request.

## Author Contributions

**Fernando de Sá Ribeiro** -Data curation; Formal analysis; Investigation; Methodology; Validation; Visualization; Writing - review & editing.

**Luís Maurício T. R. Lima -** Conceptualization; Data curation; Formal analysis; Funding acquisition; Investigation; Methodology; Project administration; Supervision; Validation; Visualization; Roles/Writing - original draft; Writing - review & editing.

