## Supporting Material for "Temperature-Resolved Crystallography Reveals Rigid-Body Dominance Over Local Flexibility in B-Factors"

[1] Laboratório de Biotecnologia Farmacêutica (pbiotech), Faculdade de Farmácia, Universidade Federal do Rio de Janeiro, Rio de Janeiro, RJ, 21941-902, Brazil.
[2] Programa de Pós-Graduação em Química Biológica, Universidade Federal do Rio de Janeiro, Rio de Janeiro, RJ, 21941-902, Brazil.
[3] Programa de Pós-Graduação em Ciências Farmacêuticas, Faculdade de Farmácia, Universidade Federal do Rio de Janeiro, Rio de Janeiro, RJ, 21941-902, Brazil.

**Running title:** Thermal-dependence of B-factor

*To whom correspondence should be addressed

**Authors Address and Contact**
- Luis Mauricio T. R. Lima – Laboratório de Biotecnologia Farmacêutica (pbiotech), Faculdade de Farmácia, Universidade Federal do Rio de Janeiro – UFRJ, CCS, Bss24, Ilha do Fundão, 21941-590, Rio de Janeiro, RJ, Brazil. Phone/Fax: (+55-21) 3938-6639 –. Social Media: @pbiotech

**AUTHOR LIST**

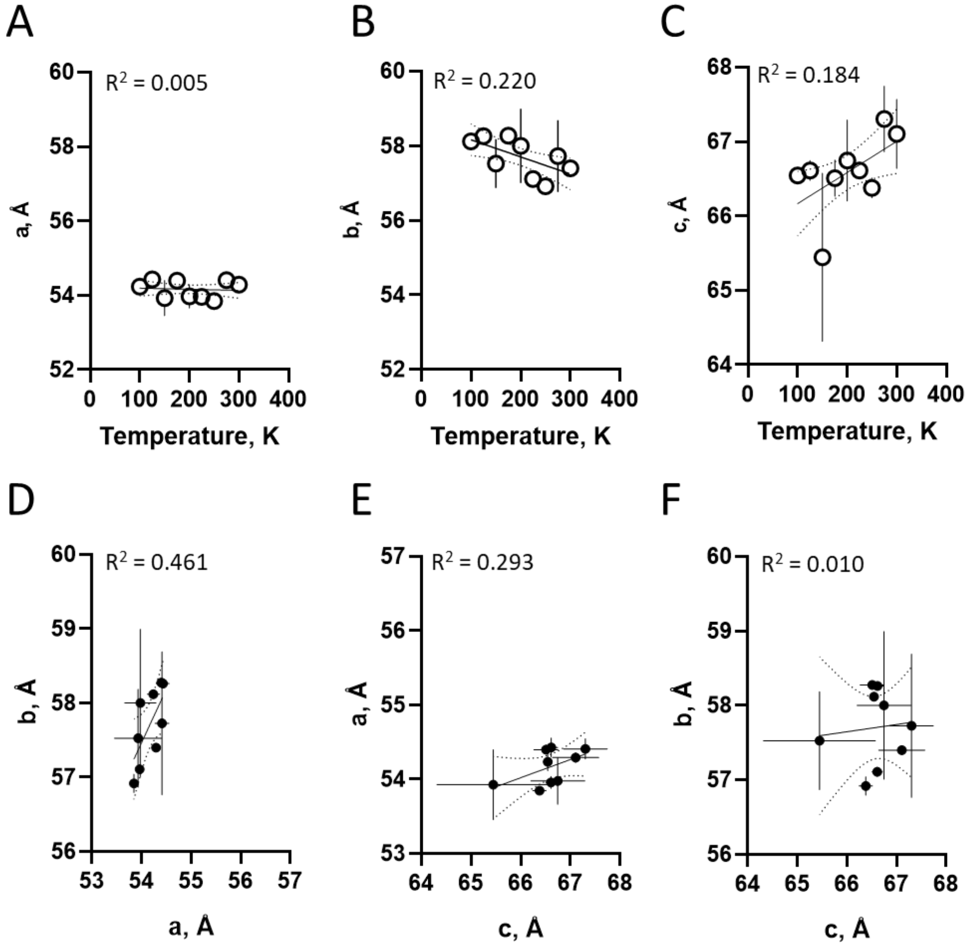


**Figure S1. Thermal-dependent changes in unit cell parameters.**

Correlation between cell unit parameters and data collection temperature for single crystal x-ray diffraction of Trypsin in $P2_12_12_1$. Continuous line are first order linear regression and dotted lines are 95 % confidence interval. The linear correlation between data collection temperature

**A)** a as a function of temperature,
**B)** b as a function of temperature,
**C)** c as a function of temperature,
**D)** b as a function of a,
**E)** a as a function of c and
**F)** b as a function of c.
Symbol is average and bar is standard deviation (n=3).

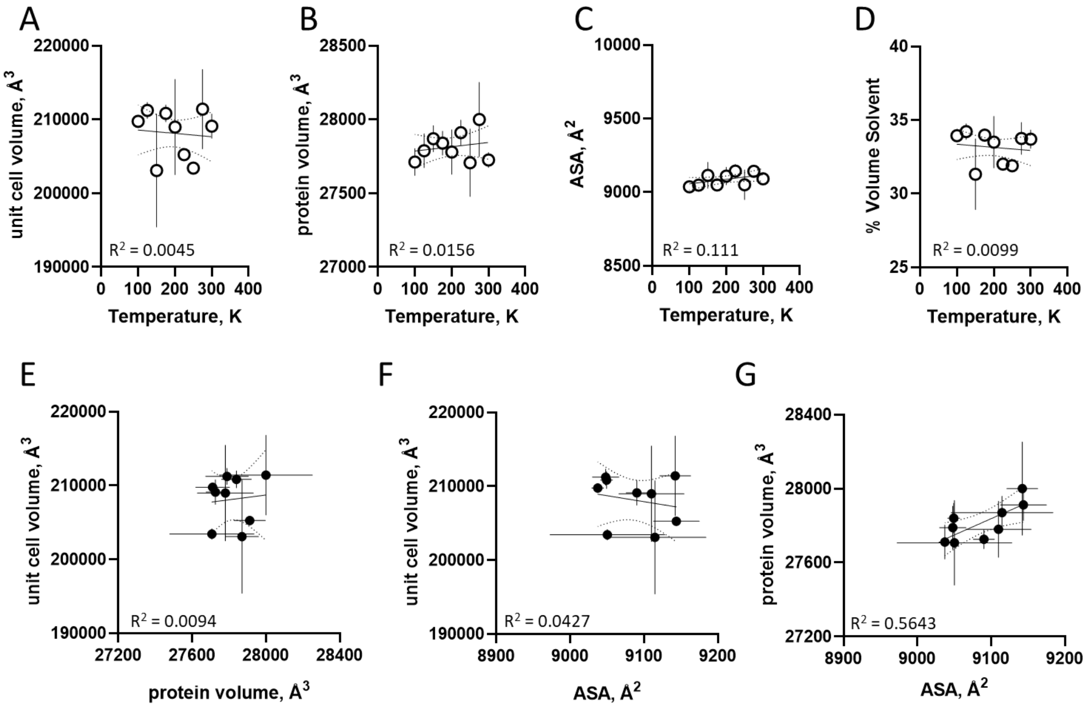


**Figure S2. Thermal-dependent changes in unit cell parameters.**

Correlation between cell unit parameters and data collection temperature for single crystal x-ray diffraction of Trypsin in $P2_12_12_1$. Continuous line are first order linear regression and dotted lines are 95 % confidence interval. The linear correlation between data collection temperature

**A)** Unit cell volume a function of temperature,
**B)** protein volume as a function of temperature,
**C)** Accessible surface area (ASA) as a function of temperature,
**D)** % volume solvent as a function of a,
**E)** Unit cell volume a as a function of protein volume,
**F)** Unit cell volume a as a function of ASA and
**G)** Protein volume a as function of ASA.
Symbol is average and bar is standard deviation (n=3).

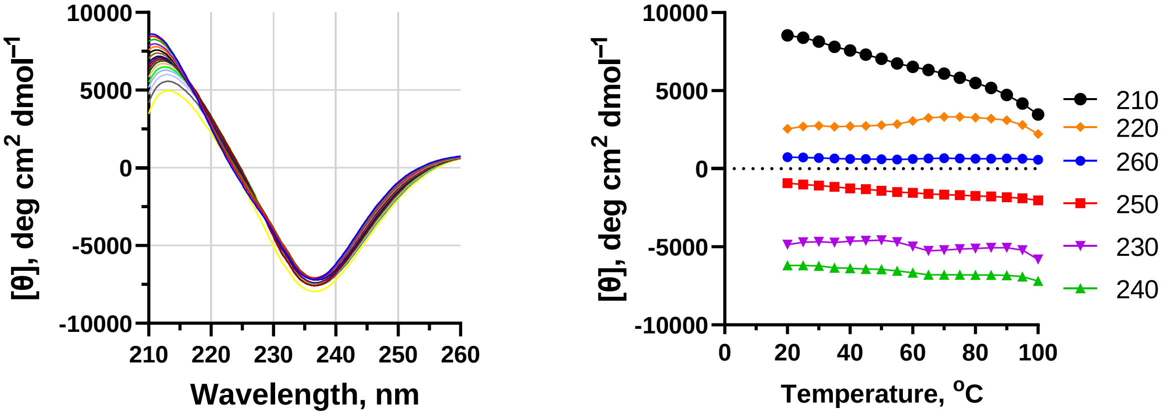


**Figure S3. Thermal-dependent conformational changes in trypsin.**
Temperature effect on lysozyme was monitored by circular dichroism providing information regarding secondary structure.
**A)** Circular dichroism spectra of trypsin at temperatures from 20 °C to 100 °C in 5 °C intervals;
**B)** Thermal curves at varying wavelengths (data from Fig. S3A; average and standard deviation from from 0.4 degree interval);
Measurements were performed with trypsin at 5 mg/mL in 20 mM dibasic potassium phosphate with benzamidine.

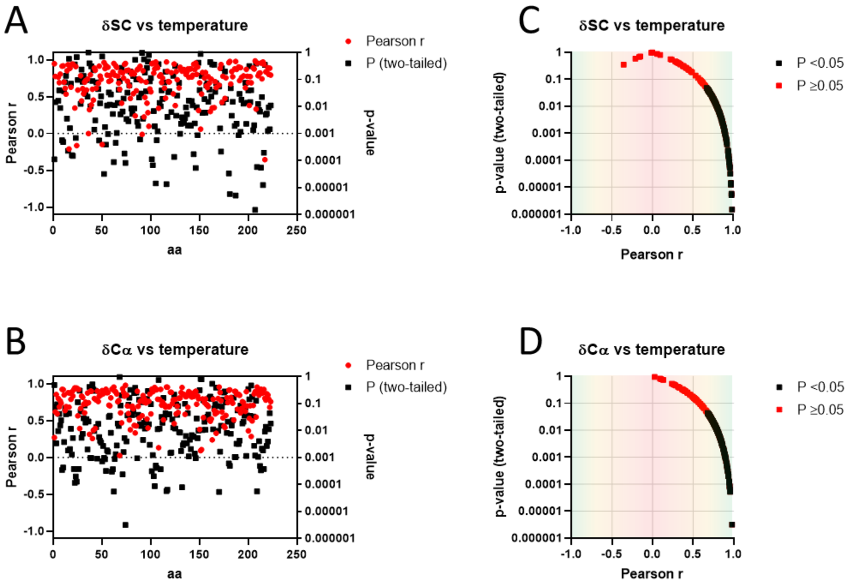


**Fig. S4. Pearson correlation between changes in trypsin conformation and temperature**

Changes in side-chain (Fig. 2) and Cα (Fig. 3) conformation were plotted as a function of temperature and analyzed for Pearson correlation.
The Pearson r and p-value (two-tailed) were plotted as a function of trypsin sequence (continuous numbering) for (A) side-chain ($r>0.67$ for $p<0.05$) and (B) Cα ($r>0.68$ for $p<0.05$).
Notice a correlation between Pearson r and p-value for both (C) side-chain and (D) Cα. Most data lie with $r>0.5$ and $p<0.05$, suggesting strong positive correlation.

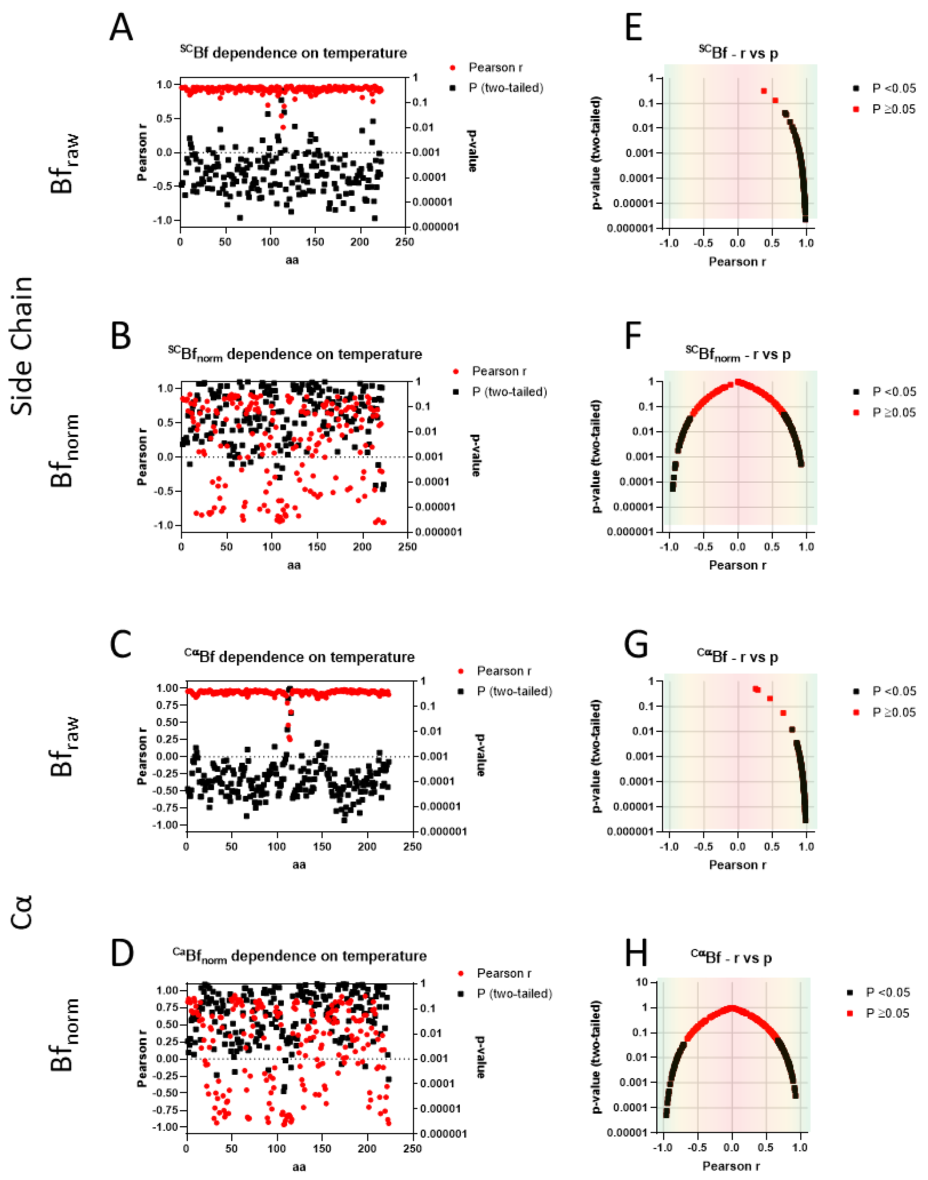


**Fig. S5. Pearson correlation between changes in trypsin B-factor and temperature**

Changes in side-chain (Fig. 5) and Cα (Fig. 6) B-factors were plotted as a function of temperature and analyzed for correlation.
The Pearson r and p-value (two-tailed) were plotted as a function of trypsin sequence (continuous numbering) for side chains (A, B) and Cα (C, D).
Notice a correlation between Pearson r and p-value for both
(E, F) side-chain ($Bf_{raw}$, $r>0.68$ for $p<0.05$; $Bf_{norm}$, $r>|0.67|$ for $p<0.05$) and
(G, H) Cα ($Bf_{raw}$, $r>0.78$ for $p<0.05$; $Bf_{norm}$, $r>|0.67|$ for $p<0.05$).
Most data for raw B-factor lies with $r>0.8$ and $p<0.05$ (E, G), suggesting strong positive correlation, while normalization (F, H) reduces correlation.

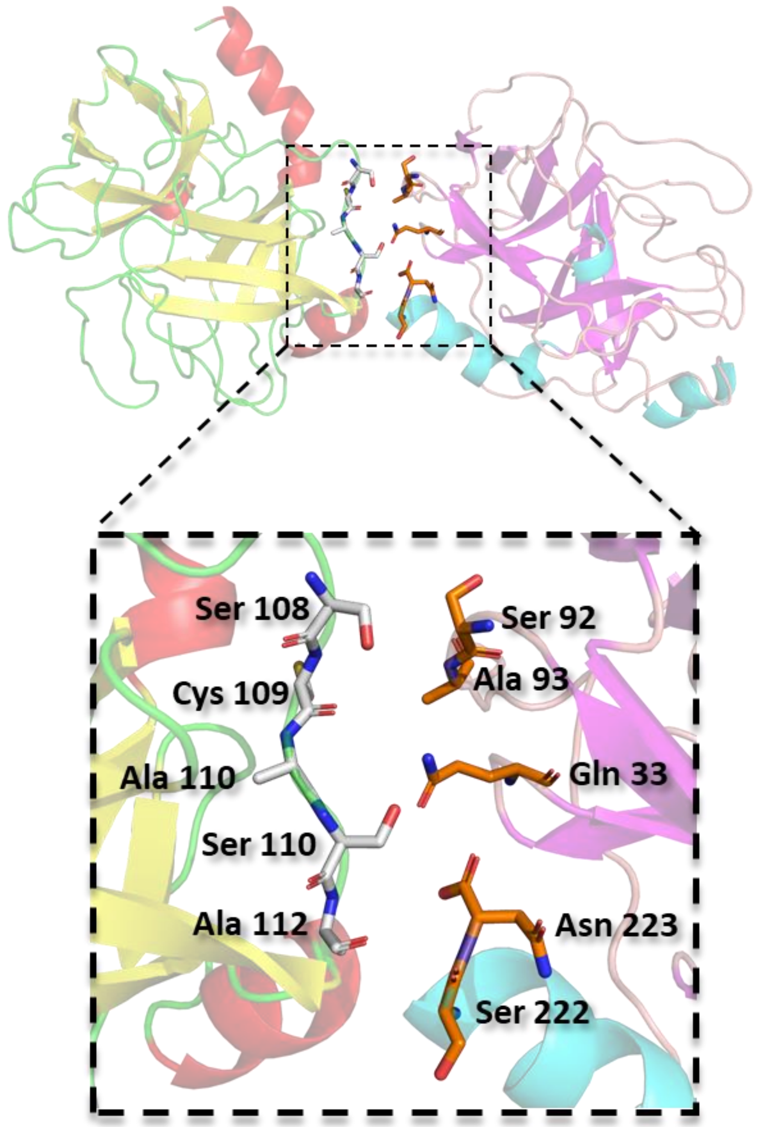


**Figure S6. Representation of the crystal contact generated by the symmetry found in the $P2_12_12_1$ space group.**
The structure of PDB: 9AVX is represented in cartoon with colors corresponding to secondary structures: yellow for alpha helices, red for beta sheets, and green for coils. The residues important for crystal contact are represented by sticks and colored according to elements: gray for carbon, blue for nitrogen, red for oxygen, and yellow for sulfur.
The symmetry-generated structure produced by PyMOL within a distance of 4 Å is represented in cartoon with colors corresponding to secondary structures: cyan for alpha helices, purple for beta sheets, and pink for coils.
The residues important for crystal contact are represented by sticks and colored according to elements: orange for carbon, blue for nitrogen, red for oxygen, and yellow for sulfur.
The residues important for the contact are labeled with their three-letter codes and respective positions.

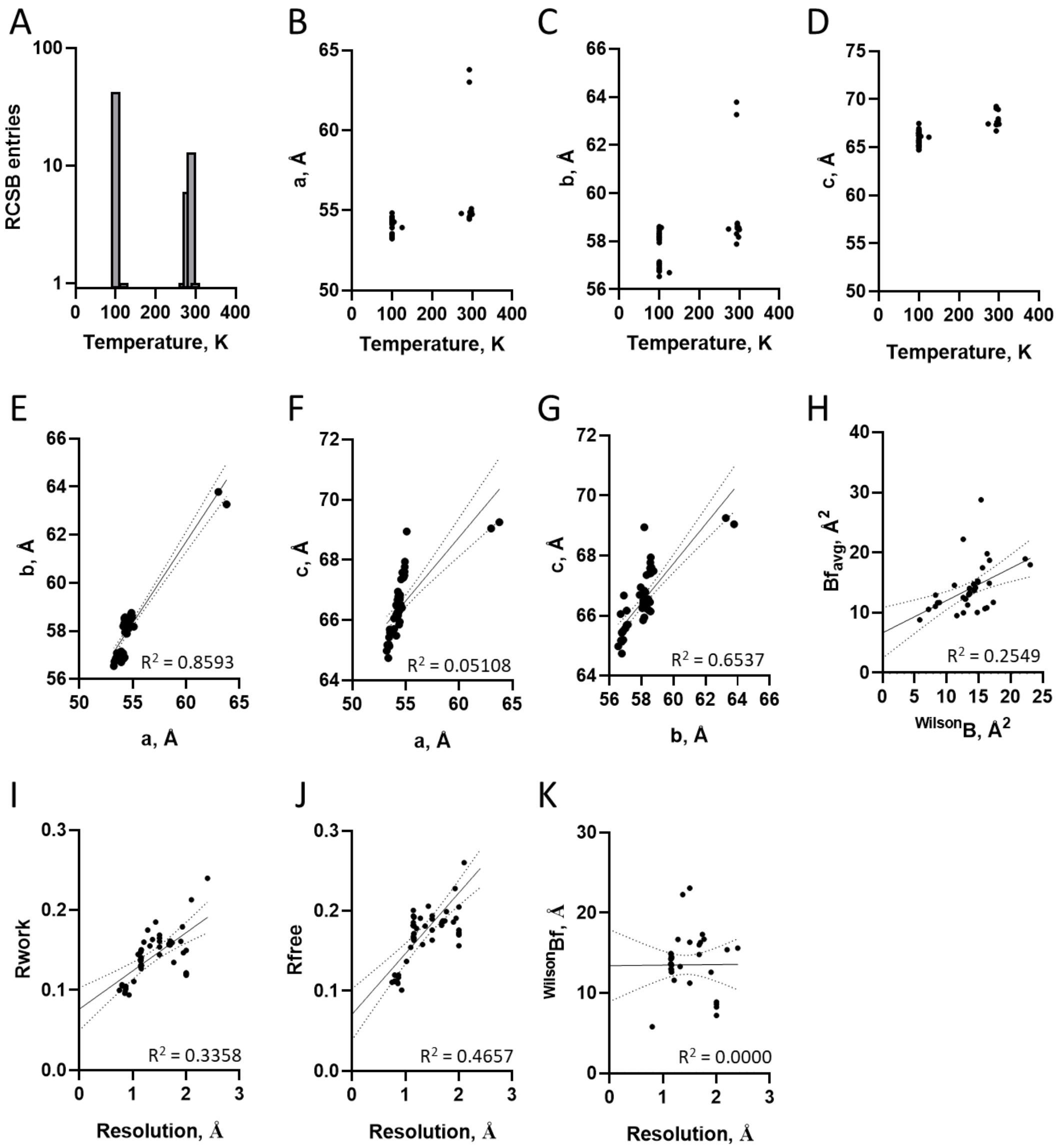


**Figure S7. Orthorhombic trypsin-benzamidine structures in the RCSB**

Data from the RCSB (access May, 2024) for trypsin-benzamidine in $P2_12_12_1$.

**A)** Distribution of entries according to data collection temperature, found between 100 k to 300 K.

**B)** Unit cell length in the a axis as a function of temperature.

**C)** Unit cell length in the b axis as a function of temperature.

**D)** Unit cell length in the c axis as a function of temperature.

Correlation between data collection temperature (from 100 k to 300 k); **(E)** unit cell length in the b axis as a function of a, **(F)** unit cell length in the b axis as a function of a **(G) u**nit cell length in the c axis as a function of b, **H)** Correlation between B-factors (all trypsin with benzamidine $P2_12_12_1$) from Wilson estimate and average from final structure model (39 entries). **(I)** distribution of Rwork and Rfree for deposited structures.

Correlation between data collection temperature (from 100 k to 300 k); **(J)** Rfree as a function of Rwork, **(K)** Rfree as a function of Wilson and **(L)** Rwork as a function of Rwork

Continuous line are first order linear regression and dotted lines are 95 % confidence interval.

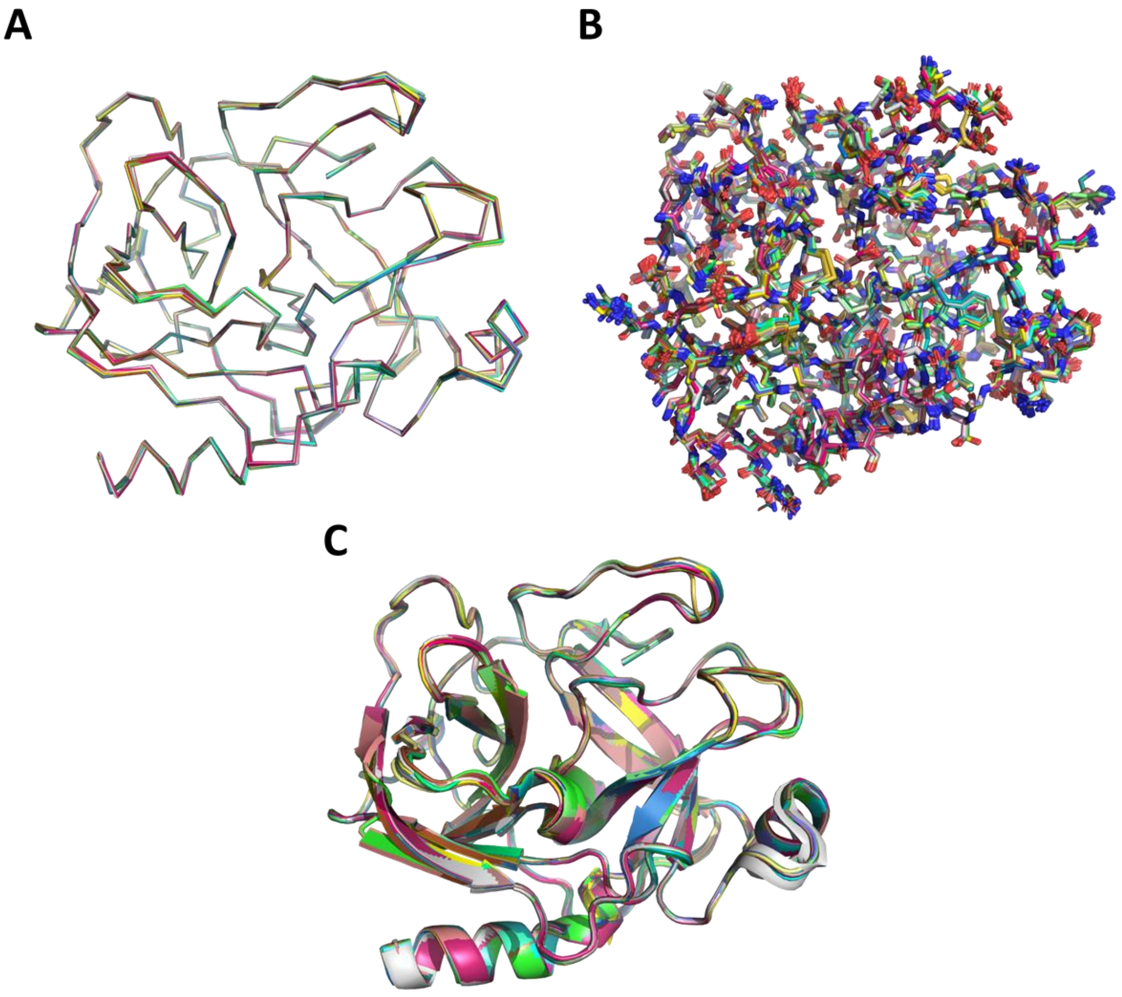

**Figure S8. Alignment of trypsin-benzamidine complexes from the RCSB. Data from the RCSB (access May, 2024) for trypsin-benzamidine in $P2_12_12_1$.**

**A)** backbone,

**B)** side chains and

**C)** cartoon
